# MotiMul: A significant discriminative sequence motif discovery algorithm with multiple testing correction

**DOI:** 10.1101/2020.08.21.261024

**Authors:** Koichi Mori, Haruka Ozaki, Tsukasa Fukunaga

## Abstract

Sequence motifs play essential roles in intermolecular interactions such as DNA-protein interactions. The discovery of novel sequence motifs is therefore crucial for revealing gene functions. Various bioinformatics tools have been developed for finding sequence motifs, but until now there has been no software based on statistical hypothesis testing with statistically sound multiple testing correction. Existing software therefore could not control for the type-1 error rates. This is because, in the sequence motif discovery problem, conventional multiple testing correction methods produce very low statistical power due to overly-strict correction. We developed MotiMul, which comprehensively finds significant sequence motifs using statistically sound multiple testing correction. Our key idea is the application of Tarone’s correction, which improves the statistical power of the hypothesis test by ignoring hypotheses that never become statistically significant. For the efficient enumeration of the significant sequence motifs, we integrated a variant of the PrefixSpan algorithm with Tarone’s correction. Simulation and empirical dataset analysis showed that MotiMul is a powerful method for finding biologically meaningful sequence motifs. The source code of MotiMul is freely available at https://github.com/ko-ichimo-ri/MotiMul.

## I. Introduction

Sequence motifs are short biological sequence patterns with biological functions, and play essential roles in intermolecular interactions. For example, the DNA sequence pattern “CACGTG” is the binding motif for the transcription factor (TF) URF1. URF1 regulates the expression of its target genes by interacting with promoter regions containing the DNA sequence pattern [1]. The discovery of novel sequence motifs is therefore an essential step in the elucidation of gene function, and the computational identification of sequence motifs has long been studied in bioinformatics.

To date, various bioinformatics tools have been developed for finding sequence motifs. Some of these tools discover sequence motifs by searching for frequently occurring sequence patterns from a single dataset consisting of a large amount of sequence data [2]–[6]. Alternatively, some tools use two sequence datasets: positive and negative datasets. They detect sequence patterns that specifically appeared in the positive datasets as the sequence motifs [7]–[11]. The latter approach is called discriminative sequence motif discovery, and is expected to improve the accuracy of the software over that of the former approach, if appropriate negative datasets are used. In this study, we focused on the development of an algorithm for discriminative sequence motif discovery.

Some methods, such as DREME [9], detect discriminative sequence motifs based on statistical hypothesis testing. Unlike machine learning-based approaches, statistical hypothesis testing has the advantage that it can control for the type-1 error rate; that is, they can keep the false positive rate below a user-defined significance level. This approach is effective for preventing the detection of invalid sequence motifs. When we apply hypothesis tests to many candidate sequence motifs, it is necessary to conduct multiple testing correction to reduce the detection of false positives. However, conventional multiple testing correction causes a significant decrease in statistical power. For example, Bonferroni’s correction method, which is widely used in biological research [12], [13], calculates an adjusted significance level by dividing the original significance level by the number of hypothesis tests. In the discriminative sequence motif discovery problem, the significance level obtained using Bonferroni’s correction is too small. Therefore, previous motif discovery tools based on statistical hypothesis testing did not include statistically sound multiple testing correction.

There have been several previous studies on sequence mining methods based on statistical hypothesis testing. Low-Kam *et al.* considered the statistical significance of frequent sub-sequences in a single dataset consisting of several sequences [14]. This study defined subsequences as ordered sequences that did not require continuity, and the null model as a model in which the probability of occurrence of the characters is independent. Jelovic *et al.* developed StatRepeats, which detects significantly numerous repeat sequences by enumerating identical contiguous sequences that appeared multiple times in the genome [15]. Nakamura *et al.* also proposed an algorithm that detects significantly frequent repeat sequences in the genome. Their methods can consider sequence patterns containing mismatches, unlike StatRepeats [16]. In recent years, Koulouras and Frith developed Nullomers Assessor, which analyzes significant absent sequences in genomes or proteomes [17]. Although these methods effectively extract biologically meaningful sequence patterns from massive amounts of sequence data, we cannot directly apply these methods to the discriminative sequence motif discovery problem.

In this study, we developed MotiMul, an algorithm that can discover significant discriminative sequence motifs with statistically sound multiple testing correction. The fundamental idea underlying the algorithm is the utilization of Tarone’s correction, which improves statistical power by ignoring hypotheses that never become statistically significant under given conditions [18]. Data mining methods for bioinformatics based on Tarone’s correction have been the subject of active research in recent years, and have been applied to itemset mining [19]–[21], graph mining [22], genomic interval analysis [23], [24], genome-wide association studies [25], [26], time-series analysis [27], survival analysis [28], and sequence analysis [17]. MotiMul can efficiently enumerate significant sequence motifs of arbitrary length by integrating Tarone’s correction with a variant of the PrefixSpan algorithm, a frequent sequence pattern mining method [29].

We first applied MotiMul to a simulated dataset, and showed that MotiMul was more sensitive than the conventional Bonferroni’s correction. Next, we evaluated basic measures of software performances, such as the statistical properties and computational speed, of the MotiMul using simulated datasets. Finally, we applied MotiMul to three empirical ChIP-seq datasets, and discovered significant sequence motifs contributing to transcriptional regulation.

## II. Methods

### A. Notation

Let Σ be an alphabet, which is defined as the set of all characters to be considered in the analysis. In this study, because we focused on the DNA sequence analysis, Σ was {*A, T, C, G*}. Let *s* be a sequence, defined as a contiguous ordered list of characters. *s* is also represented by *s*_1_*s*_2_…*s*_*L*_, where *s*_*i*_ ∈ Σ for 1 ≤ *i* ≤ *L*. Here, we call *L* the sequence length. A sequence *s* is called a subsequence of another sequence *t*, if there is a positive integer *i* that satisfies *s*_1_ = *t*_*i*_, *s*_2_ = *t*_*i+*1_, …, *s*_*L*_ = *t*_*i*+*L*−1_ where *L* is the sequence length of *s*. In this case, *t* is also called a supersequence of *s*. Additionally, we define * as the wildcard character, which represents any one of the characters in the alphabet.

Let *D*_*p*_ and *D*_*n*_ be the positive and negative datasets, respectively. We define *n*_*p*_ and *n*_*n*_ as the number of sequences included in the positive and negative datasets, respectively. We also define *N* as *n*_*p*_ + *n*_*n*_.

### B. Hypothesis testing of sequence patterns

We detected discriminative sequence motifs based on statistical hypothesis testing. We envisaged a sequence pattern with the wildcard less than or equal to the user-defined parameter *w* as a candidate sequence motif. For a candidate sequence motif *s*, the null hypothesis was that the frequency with which a sequence in the datasets is a supersequence of *s* is independent of the differences in the datasets. If the null hypothesis was rejected by the one-sided Fisher’s exact test, we accepted the candidate sequence motif *s* as a sequence motif.

Let *x* and *y* be the numbers of the sequences that are supersequences of *s* in *D*_*p*_ and *D*_*n*_, respectively. In the Fisher’s exact test, the contingency table is represented as follows:

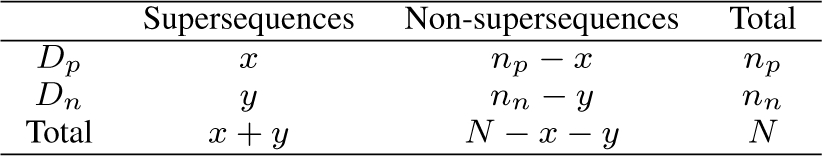

Here, because the null distribution follows the hypergeometric distribution, the *p*-value is calculated as follows:

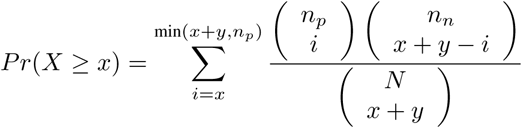

If the *p*-value is smaller than the user-defined significance level *α*, the null hypothesis is rejected. In this study, we set *α* to 0.05 in all experiments.

### C. Multiple testing correction

To exhaustively discover significant sequence motifs, we need to perform hypothesis tests on all candidate sequence motifs. If all the sequence lengths in the dataset are *L* and the candidate sequence motifs are of arbitrary lengths, the number of hypotheses is 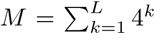 when the parameter *w* is 0. When we conduct multiple hypothesis tests simultaneously, we must adjust the significance level to prevent detecting many false-positives. In this study, we adjusted the significance level so that the family-wise error rate (FWER) was controlled. The FWER is the probability of detecting at least one false positive when testing multiple hypotheses. Bonferroni’s correction is a conventional method for controlling FWER, producing an adjusted significance level by dividing the original significance level *α* by the number of hypothesis tests *M*. However, Bonferroni’s correction has the problem that the statistical power becomes too low with a large number of hypotheses, because the significance level becomes very small. Therefore, the application of Bonferroni’s correction to the discriminative sequence motif discovery problem should be ineffective.

Tarone’s correction is a technique for improving the statistical power in multiple testing correction for controlling the FWER [18]. This method first divides all statistical hypotheses into untestable and testable hypotheses, and calculates the adjusted significance level by dividing the original significance level *α* by the number of testable hypotheses only. Here, a testable hypothesis is a hypothesis that can become statistically significant, and an untestable hypothesis is a hypothesis that never becomes statistically significant under the given conditions. In most cases, because the number of testable hypotheses is much less than the number of all hypotheses, Tarone’s correction gives a much higher significance level and statistical power than Bonferroni’s correction.

We explain the concept of untestable and testable hypotheses using the aforementioned contingency table in the Fisher’s exact test. When the one-sided Fisher’s exact test is performed and the values of *x* + *y* and *n*_*p*_ are given, a larger *x* leads to a smaller *p*-value of the hypothesis test. The maximum possible value of *x* is min(*x* + *y*, *n*_*p*_). We can then obtain the lower bound of the *p*-value based on the maximum possible value of *x* before the hypothesis test is performed. If the lower bound of the *p*-value is larger than the significance level, the hypothesis test never becomes statistically significant; that is, it is an untestable hypothesis. Otherwise, the hypothesis is testable.

The detailed procedure of the general Tarone’s correction is as follows. We first calculate the lower bounds of the *p*-values of all hypothesis tests under the given conditions, and sort the hypotheses in the ascending order of the lower bounds of the *p*-values. Here, if the lower bound of the *p*-value of the (*k* + 1)-th hypothesis is larger than 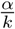, subsequent hypotheses are untestable hypotheses. Therefore, we search for the minimum *k* satisfying this condition and use 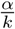 as the adjusted significance level.

### D. Enumeration of significant discriminative sequence motifs

Tarone’s correction requires the enumeration of the testable hypotheses, but direct enumeration is computationally intractable in discriminative sequence motif discovery due to the massive number of the hypotheses. The development of an efficient method of enumeration is a computational challenge in this problem. In recent years, the efficient enumeration methods based on the frequent pattern mining algorithm have been proposed for significant itemset [19] and subgraph [22] detection. Following these studies, we enumerated the testable hypotheses using a variant of the PrefixSpan algorithm, a frequent sequence pattern mining method [29]. The variant of the PrefixSpan algorithm can discover all candidate sequence motifs whose frequency of supersequences in the database is higher than the user-defined value λ.

We defined *l*(λ) as the lower bound of *p*-value when λ = *x* + *y* in the above contingency table, and *f* (λ) as follows:

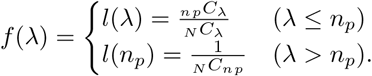

Here, *l*(λ) > *l*(*n*_*p*_) when λ > *n*_*p*_, and *f*(λ) is a monotonically decreasing function. Additionally, let *I*_λ_ be the set of candidate sequence motifs enumerated by the variant of the PrefixSpan algorithm from both *D*_*p*_ and *D*_*n*_ when the user-defined threshold is λ, and *m*_*λ*_ be |*I*_λ_|. Then, the number of the testable hypotheses is calculated as *m*_*λ*_ satisfying the two following conditions: *f* (λ − 1)*m*_λ_ > *α* and *f*(λ)*m*_*λ*+1_ ≤ *α*, and the adjusted significance level becomes *α*/*m*_*λ*_ [19], [22]. We defined *λ\** as λ satisfying this condition.

We developed two approaches to obtain *λ\**. The first approach was a decremental search algorithm. In this method, we first set λ to *N*, which is the maximum possible frequency, and repeated the application of the variant of the PrefixSpan algorithm to the datasets while lowering the threshold λ one by one. When we found a λ satisfying the condition *f*(λ – 1)*m_λ_ > α* for the first time, we set *λ\** to the λ. The search method was firstly proposed as a part of the LAMP algorithm for finding significant itemsets [19]. The second method was an incremental search algorithm. In this method, in contrast to the decremental search, we first set λ to 1 and conducted the variant of the PrefixSpan algorithm repeatedly while increasing the threshold λ one by one. When we discovered a λ satisfying the condition *f*(λ)*m*_*λ*_+1 ≤ *α* for the first time, we set *λ\** to the λ. Note that we did not have to run the variant of the PrefixSpan algorithm for λ satisfying *f*(λ)> *α* when λ is small. This is because *m*_λ+1_ ≥ 1 and thus *f*(λ)*m*_λ+1_ > *α* when *f*(λ) > *α* regardless of the value of *m*_λ+1_. This search method was first proposed for finding the significant subgraphs [22]. We implemented these algorithms using C++ as a program named MotiMul. The source code is freely available at https://github.com/ko-ichimo-ri/MotiMul.

### E. Dataset preparation

To investigate the statistical power and the basic performance of MotiMul, we generated simulated datasets and applied MotiMul to the datasets. Although we created different simulated datasets for separate investigations, we used the following three characteristics for the generation of all simulation datasets. First, a simulated dataset consisted of a positive and a negative dataset, and the number of the sequences in the positive dataset and that in the negative dataset were the same. Second, the sequence lengths of the sequences in the dataset were all the same. Third, characters other than the artificial sequence motif were determined randomly with equal probabilities from the characters in Σ.

For the empirical dataset analysis, we applied MotiMul to ChIP-seq datasets for three TFs from K562 cells: USF1, GABP, and SRF. We obtained the ChIP-Seq peak data from the ENCODE project [30], and we used the regions within the range of ±50 base pairs from the peak as the positive dataset. There were 24059 sequences in the USF1 dataset, 12559 in the GABP dataset, and 5453 in the SRF dataset. We generated the negative dataset by shuffling each sequence in the positive dataset, while preserving the dinucleotide frequencies. We used the ushuffle software for this step [31].

## III. Results

### A. Comparison analysis between Bonferroni’s correction and MotiMul

We first investigated the difference in statistical power between Bonferroni’s correction and MotiMul based on Tarone’s correction, using a simulated dataset. We used a simulation dataset in which the number of sequences for each dataset was 1000 and the sequence length was 100. Additionally, we considered an 8-character string “TTATGCAA” as an artificial sequence motif, and embedded the sequence motif in 100 sequences in the positive dataset.

We applied Bonferroni’s correction and MotiMul to this simulated dataset. When we used Bonferroni’s correction, the number of hypothesis tests was 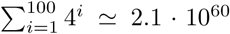 and the adjusted significance level was 2.3 · 10^−62^. Because the adjusted significance level was too low, we could not detect the sequence motifs. When we used the MotiMul with the parameter *w* = 0, the number of hypothesis tests was 83151 and the adjusted significance level was 6.0 · 10^−7^. The significance level was much larger than that of Bonferroni’s correction, and thus MotiMul could discover five significant sequence motifs: “TTATGCAA” (*p*-value: 6.1·10^−21^), “TTAT-GCA” (1.4 10^−12^), “TATGCAA” (2.5·10^−9^), “TTATGCAAA” (3.4 · 10^−8^) and “TTTATGCAAA” (5.0 · 10^−8^). The sequence with the lowest *p*-value was the true sequence motif, and all other significant sequence motifs were subsequences or supersequences of the true sequence motif. These results showed that MotiMul is effective for finding significant discriminative sequence motifs using statistically sound multiple testing correction.

### B. Investigation of the basic performances of MotiMul

We investigated the basic performance of MotiMul by applying it to simulated datasets. We first evaluated whether MotiMul can control the FWER. In this analysis, we used a simulated dataset in which the number of sequences for each dataset was 100 and the sequence length was 100. We did not embed the artificial sequence motif in the positive dataset. Then, we created 1000 simulation datasets and applied MotiMul with the parameter *w* = 0 to all datasets. We detected at least one motif in 20 datasets; that is, the empirical FWER was 0.02. This result shows that MotiMul can appropriately control the FWER.

We then examined the dependence of the computational speed of MotiMul on the sequence lengths and the dataset sizes. We compared the computational speed of MotiMul based on incremental search with that based on decremental search. For these analyses, we considered “TTATGCAA” to be an artificial sequence motif, and we embedded the sequence motif in all sequences in the positive dataset. For the analysis of the dependence on the sequence lengths, we used a simulated dataset in which the number of sequences in each dataset was 100. We also used ten sequence lengths between 20 and 200 with 20 increments. For the analysis of the dependence on the dataset sizes, we used a simulation dataset in which the sequence lengths were 100. We used ten number of sequences for each dataset between 25 and 250 with increments of 25. We created 100 simulated datasets for each sequence length and each number of sequences, and calculated the averaged execution time. We set the parameter *w* of MotiMul to 0 in these analyses. The computation was performed on an Intel(R) Core(TM) i7-8650U 1.90GHz CPU with 16GB of memory.

Fig. 1 shows the way in which the execution time depends on the sequence lengths and dataset sizes. In both analyses, the execution time monotonically increased with increasing sequence lengths and increasing number of sequences. We verified that MotiMul using incremental search was faster than MotiMul using decremental search in all cases. The difference was more pronounced when the dataset size was large. While the running time ratio of the incremental search and the decremental search was 3.93 when the dataset size was 25, it was 23.87 when the dataset size was 250. However, the running time ratio was almost constant when for different sequence lengths. Our finding that the incremental search was more effective than the decremental search is consistent with previous studies for significant itemset mining [32] and significant subgraph mining [22].

**Fig. 1.**
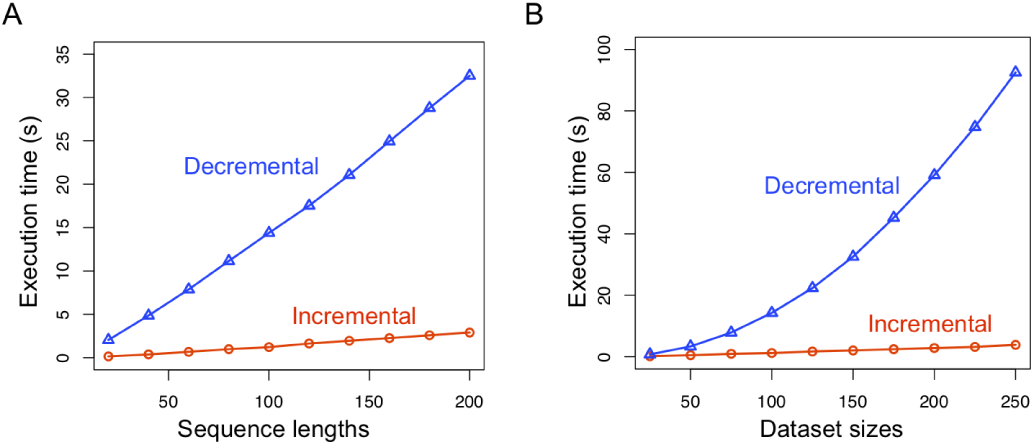
Execution times of incremental search and decremental search when (A) the sequence lengths or (B) the dataset sizes were changed. The x-axes represent the sequence length or the dataset size, and the y-axes represent the execution times. Incremental search and decremental search are represented by the red and blue lines, respectively.

We next evaluated the dependence of the computational speed and the number of testable hypotheses on the user-defined parameter *w*, the maximum value of the wildcard character in a sequence motif. We used the same datasets as those investigated for differences in computational speed due to differences in the sequence lengths and the dataset sizes. The computation was performed on an Intel Xeon Gold 6148 3.7GHz CPU with 154GB of memory. Fig. 2 shows the results. The execution time and the number of testable hypotheses increased with increasing value of the parameter *w*.

**Fig. 2.**
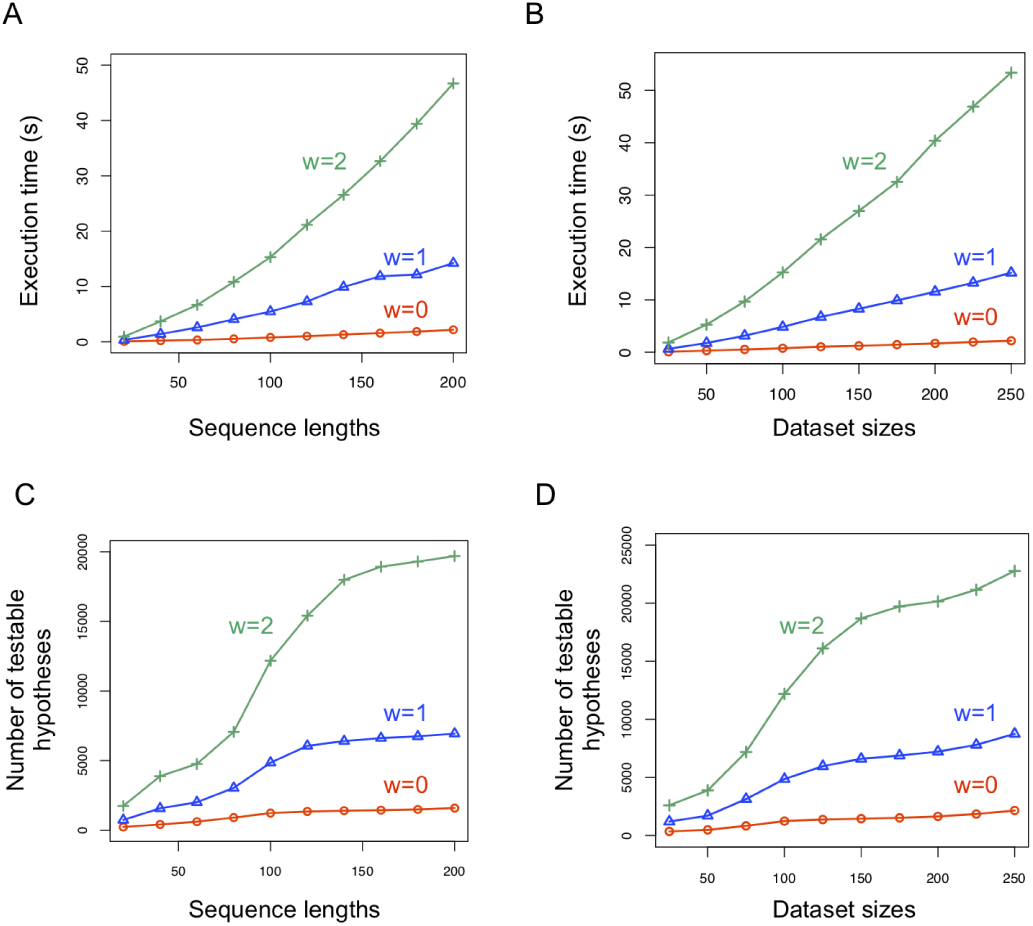
Dependence of execution times and number of testable hypotheses on the parameter *w*. Panels (A) and (C) represent the results when the sequence lengths were changed, and panels (B) and (D) represent the results when the dataset sizes were changed. Additionally, panels (A) and (B) represent the execution times, and panels (C) and (D) represent the number of the testable hypotheses. The x-axes represent the sequence lengths or the dataset sizes, and the y-axes represent the execution times or the number of testable hypotheses. The red, blue, and green lines represent the parameter values *w* = 0, *w* = 1 and *w* = 2, respectively.

### C. Theoretical analysis of the upper bound of λ*

We theoretically analyzed the upper bound of *λ*\* when the parameter *w* = 0. We first estimated the upper bound of *m*_*λ*\*_. Because a sequence has at most _*L*_*H*_2_ subsequences, the number of all subsequences in the total dataset is at most *N*_L_H_2_. Therefore, 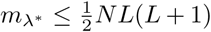. We next investigated the condition that satisfied λ* > *n*_*p*_. From the conditions met by *λ*\*, *α* < *f*(λ * 1)*m*_*λ*\*_ has to be met. Therefore, 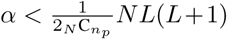 must be satisfied. Because this formula is not satisfied unless *L* is very large compared with *N*, we do not have to assume this situation in general sequence motif analyses. Therefore, we can assume λ* ≤ *n*_*p*_.

Here, the following inequality about *f*(*x*) holds:

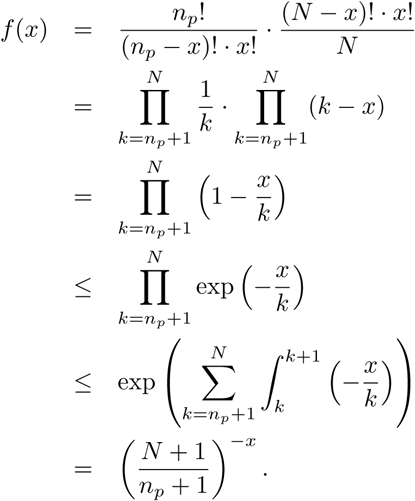

Accordingly, when 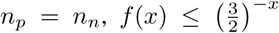. From the conditions met by *λ*\*, 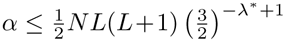
. Therefore, the upper bound of *λ*\* is calculated as the following formula:

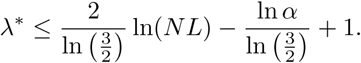

That is, *λ*\* = *O*(ln(*NL*)). This result indicates that incremental search is more effective than decremental search, because *λ*\* should be quite small. Note that the same discussion holds when the dataset size of the positive dataset is a constant multiple of the dataset size of the negative dataset. Although we set the parameter *w* to 1 or more, a similar discussion holds by changing only the upper bound of *m*_*λ*\*_.

### D. ChIP-seq data analysis

Finally, we detected significant TF-binding sequence motifs using three ChIP-seq datasets. We used MotiMul with the incremental search algorithm. The computation was performed on an Intel Xeon Gold 6148 3.7GHz CPU with 154GB of memory. Table 1 summarizes the execution results. MotiMul produced sufficient significance levels and could find significant sequence motifs in a reasonable time except when we applied MotiMul with the parameter *w* = 2 to the USF1 dataset. In this case, MotiMul could not return the results due to a shortage of computational memory. In all cases in which we could experiment, λ* became small values.

**TABLE I.** Summarization of the execution results

|  | USF1 |  |  | GABP |  |  | SRF |  |  |
| --- | --- | --- | --- | --- | --- | --- | --- | --- | --- |
| The parameter w | 0 | 1 | 2 | 0 | 1 | 2 | 0 | 1 | 2 |
| The number of testable hypothesis | 161696 | 1173116 | - | 87988 | 596270 | 2396758 | 41632 | 262562 | 1005961 |
| The adjusted significance level | $3.1 \cdot 10^{-7}$ | $4.3 \cdot 10^{-8}$ | - | $5.7 \cdot 10^{-7}$ | $8.4 \cdot 10^{-8}$ | $2.1 \cdot 10^{-8}$ | $1.2 \cdot 10^{-6}$ | $1.9 \cdot 10^{-7}$ | $5.0 \cdot 10^{-8}$ |
| The number of rejected hypothesis | 7656 | 53257 | - | 2287 | 11070 | 30421 | 2544 | 18939 | 95880 |
| The execution time (s) | 1130.84 | 8476.32 | - | 371.09 | 3186.31 | 13414.48 | 101.92 | 1004.19 | 5109.23 |
| $\lambda^*$ | 22 | 25 | - | 21 | 24 | 26 | 20 | 23 | 25 |

We compared the sequence motifs detected by MotiMul with those detected by MOCCS [5]. MOCCS identifies sequence motifs of the length *k* by quantifying the shape of the relative frequency distribution of each *k*-mer around the peak regions as an AUC score. MOCCS requires the user-defined parameter *k* and cannot enumerate sequence motifs of arbitrary length. We used *k* = 8 as in the original MOCCS paper.

Table 2–4 shows the lists of five sequence motifs with the low *p*-values detected by MotiMul and those with the high AUC scores identified by MOCCS for each TF. We confirmed that the results detected by MotiMul with the parameter *w* = 0 and those of MOCCS were almost identical. For example, in the USF dataset analysis, MotiMul discovered “CACGTG” as the sequence motif with the lowest *p*-value, and MOCCS identified “GTCACGTG”, which is a supersequence of “CACGTG”, as the sequence motif with the highest AUC score. In the SRF dataset analysis, we could not find the sequence motif identified by MOCCS corresponding to “GGAAGT”, which was the lowest *p*-value detected by MotiMul, at a glance. However, “GGAAGT” is a subsequence of the complementary sequence of “ACTTCCGG”, which had the fifth-highest AUC score identified by MOCCS, and thus we could conclude that the results were consistent. At present, MotiMul cannot treat the complementary sequences as the same sequences. This limitation is not a problem in the analysis of RNA sequences or protein sequences, but it may cause misinterpretation in DNA sequence analysis, as in this case.

**TABLE II.** Detected sequence motifs for the USF1 dataset

| rank | MotiMul(w=0) | MotiMul(w=1) | MOCCS |
| --- | --- | --- | --- |
| 1 | CACGTG | CACGTG | GTCACGTG |
| 2 | CACGT | CAC*TG | GTCACATG |
| 3 | ACGTG | CACGT | GTCAGGTG |
| 4 | ACGT | CA*GTG | TCAGCTGA |
| 5 | CACG | ACGTG | ACTTCCGG |

**TABLE III.**
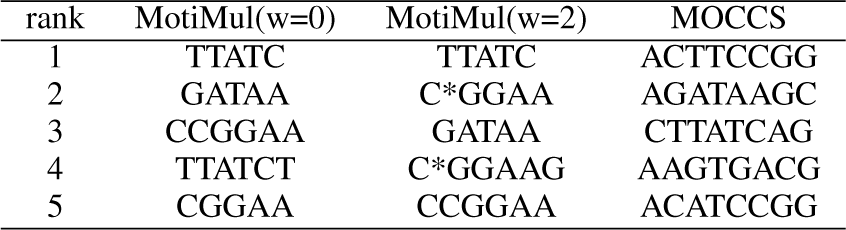
Detected sequence motifs for the GABP dataset

**TABLE IV.**
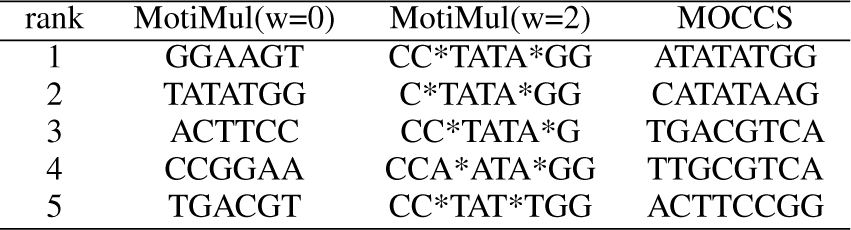
Detected sequence motifs for the SRF dataset

When the parameter *w* was set to 1, MotiMul found the “CAC*TG” and “CA*GTG” sequence motifs in the USF1 dataset analysis. These sequence motifs were the wildcard-masked sequences of “CACGTG”, which was the lowest *p*-value detected by MotiMul. These results were consistent with previous experimental researches which found that USF1 can bind the non-canonical sequence motifs “CAGGTG” (or the complementary sequence motif “CACCTG”) in addition to the canonical sequence motif “CACGTG” [33]. MOCCS also detected these non-canonical sequence motifs, but the sequence motifs were not represented in the JASPAR database, a standard sequence motif database [1]. When the parameter *w* was set to 2, MotiMul discovered “CC*TATA*GG” sequence motifs in the SRF dataset analysis. SRF is known to bind CC(A/T)_6_GG, which is called the CarG-box motif [34], and the bases immediately before C and after G are particularly susceptible to change. The results of our analysis were consistent with these characteristics. The sequence motifs included in the JASPAR database represented these characteristics, but the analysis results of MOCCS did not clearly show the characteristics. This is because MOCCS requires the user to specify the parameter *k*. These results indicated that MotiMul is a powerful approach for detecting biologically meaningful sequence motifs.

## IV. Discussion

In this study, we developed MotiMul, a novel method for discriminative sequence motif discovery with statistically sound multiple testing correction. MotiMul uses Tarone’s correction to adjust the significance level and discovers significant sequence motifs using a variant of the PrefixSpan algorithm. We showed that MotiMul based on incremental search could efficiently enumerate significant sequence motifs in theoretical and empirical analyses. Additionally, we verified that MotiMul effectively detected biologically meaningful sequence motifs in the empirical ChIP-seq datasets.

We envision three future extensions of MotiMul. The first is the summarization of significant sequence motifs for ease of interpretation. MotiMul can comprehensively enumerate significant sequence motifs, but the interpretation of all results is difficult because of the high number of hits detected. The main reason for the considerable increase in the number of discoveries is that many subsequences and supersequences are detected for a sequence motif. In recent years, Bellodi *et al.* proposed a technique for summarizing significant subgraphs to produce an interpretable representation [35]. This method learned compactly summarized structures of the significant subgraphs using the SLIPCOVER algorithm [36], a probabilistic logic programming method. The structure is represented as a logic program with annotated disjunctions (LPAD). This study showed that LPAD is a highly interpretable representation, which maintains discriminating power. The integration of this method with MotiMul should increase the usefulness of the software.

The second extension involves considering the number of sequence motifs in the sequences. Although in this study we only considered the presence or absence of the sequence motifs, the number of sequence motifs in a promoter region also affects gene regulation. The larger the number of the sequence motifs in the promoter region, the easier it is for the TF to bind. MotiMul may detect significant differences in the number of sequence motifs by modifying the statistical test and the search algorithm.

The third potential extension is the development of detection methods for significant sequence motif combinations. Because some genes are regulated by multiple TFs, the discovery of sequence motif combinations is an essential task for revealing the gene regulatory mechanisms [19], [37]. We should be able to detect discriminative significant sequence motif combinations by improving the search algorithm of MotiMul.

## Acknowledgments

The authors thank Mr. Yukihiro Okamoto for his helpful discussions. Computations in this research were performed using the supercomputing facilities at the National Institute of Genetics in Research Organization of Information and Systems.

